# On non-random mating, adaptive evolution and information theory

**DOI:** 10.1101/2024.11.18.624188

**Authors:** Antonio Carvajal-Rodríguez

## Abstract

Population genetics describes evolutionary processes, focusing on the variation within and between species and the forces shaping this diversity. Evolution reflects information accumulated in genomes, enhancing organisms’ adaptation to their environment.

In this paper, we propose a model that begins with the distribution of matings based on mutual fitness and progresses to viable adult genotype distribution. At each stage the changes result in different measures of information. The evolutionary dynamics of each stage of the model correspond to aspects such as the type of mating, the distributions of genotypes within matings, and the distribution of genotypes and haplotypes in the next generation. Changes in these distributions are caused by variation in fitness and result in Jeffreys divergence values other than zero. As an example, a model of hybrid sterility is developed at a biallelic locus, comparing the information indices associated with each stage of the evolutionary process.

In conclusion, the informational perspective seems to facilitate the connection of causes and effects and allows the development of statistics to contrast null models of zero information (random mating, no selection, etc.). The informational perspective could contribute to clarify, deepen and expand the mathematical foundations of evolutionary theory.

**SIMPLE SUMMARY:** The evolutionary process can be seen as a process of acquisition, storage, and updating of information by a population about the environment in which it lives. In this paper I propose a model that starts with the distribution of matings that occur according to mutual mating fitness and ends with the distribution of viable adult genotypes obtained after these matings. The result of the evolutionary dynamics associated with each stage of the model can be described in terms of information. This informational description facilitates the connection between causes and effects, as well as the development of statistics to test the null model of zero information, i.e. random mating and/or no effect of natural selection. Incorporating the informational perspective into the mathematical formalism of population genetics/genomics contributes to clarifying, expanding and deepening the mathematical description of evolutionary theory.

## 1. INTRODUCTION

The evolutionary process involves the accumulation of information in genomes, which allows living beings to improve their adaptation to the environment. We can trace the mathematical treatment of information back to Shannon [1], who, building on previous work by Nyquist [2] and Hartley [3] on communication and information, published his mathematical theory of communication in 1948, which soon expanded and became useful across a wide range of fields. In biology, there is an abundance of studies showing different descriptions and uses of information theory in a biological context, see for example [4–6]. In population genetics, one of the first authors to consider natural selection as a process of obtaining information about the environment during adaptive evolution is Kimura [7]. Spetner [8] also discusses the transmission of information during adaptive evolution, and later Maynard Smith [9] and Szathmáry [10] recognize the potential relevance of the concept of information in evolutionary theory. Mac Kay ([11]) uses this formalism to show that recombination allows the acquisition of information due to natural selection at a faster rate than parthenogenesis. In the new century, Adami demonstrates the connection between information theory and the evolution of biological complexity and molecular biology [12–14]. In 2001, Frieden et al. [15] poses evolutionary dynamics as a process that maximizes information and Strelioff et al [16] describe the mutual information between loci that captures the epistasis. However, it seems to be Frank [17–19] who first shows the equivalence between the description in terms of information and the classical dynamic description of population genetics. More recently, other authors have developed and deepened similar ideas [20–22].

Focusing on the selective component of the evolutionary process, the classical dynamic description starts with initial frequencies, on which the force of natural selection acts, allowing us to predict the final frequencies. However, given the significant difficulty in directly measuring the effect of natural selection, the informational approach becomes relevant, where we consider the fluctuation between the frequencies of the ancestral population and the new one, to obtain the information associated with the natural selection process that has caused that fluctuation [17,20]. Thus, we can obtain information about natural selection from the observed frequencies in two separate population groups, either in space or in time.

In summary, population genetics is the formal theory of evolution, focusing on the concepts of natural selection, genetic drift, mutation, recombination, etc., and the relationship of these concepts with adaptation to the environment and their effect on genetic diversity. However, in recent years a reformulation of the evolutionary process in terms of information is gaining ground.

This information-based approach has recently been extended to calculate information gained by comparing the distribution of phenotypes in matings with their distribution in the population. This has allowed for the derivation of sexual selection and assortative mating statistics, as well as the design of a hierarchy of models that enables the application of multi-model selection techniques to the study of non-random mating [23–25].

To develop this description in terms of information, the connection between Price’s equation, which describes the mean phenotypic change due to the effect of selection, and Jeffreys divergence was utilized, as established in Frank [18].

In this paper we propose a model that starts with the distribution of matings that occur according to mutual mating fitness and ends with the distribution of viable adult genotypes obtained after these matings. The evolutionary dynamics associated with each stage of the model can be described in terms of information.

The structure of the paper is as follows: first, we will briefly review Price’s equation and the notions of entropy and information. Second, we will review recent results related to mating fitness and non-random mating to show various connections with some measures and properties of information theory. Third, we propose the inclusion of the mating fitness concept in a more general model that would incorporate the genotype-phenotype map and other elements of the life cycle, such as viability. We will show that the mean change in fitness, caused by the shift in genotypic and haplotypic distributions due to sexual and natural selection, can be expressed as Jeffreys divergence. We will explicitly calculate the information for the simple case of a biallelic locus with non-random mating, where we demonstrate the different information measures obtained for a model of hybrid sterility. Finally, we will briefly comment, in light of the above and recent literature, whether the concept of information acquisition could serve as an alternative to biological fitness as a design principle and central axis of evolutionary dynamics.

## 2. PRICE EQUATION

Price’s equation is a mathematical identity that, in its most general form, can be applied to any characteristic that classifies two sets of entities separated in time and/or space [18,19,26–29]. Furthermore, Price’s equation has been considered a fundamental, unifying, and model-generating theorem in evolutionary theory [30–32]. Indeed, key theorems of evolution, such as the gene selection theorem (average excess), the fundamental theorem of phenotypic selection (Robertson’s equation), Breeder’s equation, and Fisher’s fundamental theorem can be derived from it [33], although regarding the latter, see the recent work by Ewens [34].

Price’s equation can be interpreted as what Kuhn [35] called symbolic generalizations or generalization sketches, and as such, it has been connected with many types of models, from which new variants of the equation can be derived to solve different types of problems [reviewed in 30]. Furthermore, Price’s equation can also be interpreted in terms of information [18], and as we have mentioned, this interpretation has recently been levereged to generate estimators of sexual selection patterns and assortative mating [23–25].

Price’s equation can be formalized in different ways; here we will follow Frank’s notation to express it as the mean change between two sets of entities for a character *z* indexed by different classes *i*,

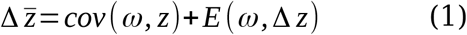

here 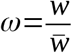.

## 3. ENTROPY AND INFORMATION

Information is concerned with the decrease in entropy or uncertainty in a system. Shannon [1] defined the uncertainty or entropy of a discrete random variable as *X*, which takes values *x*_i_ with probability *p*_i_, as

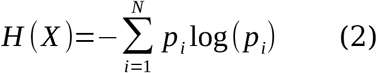

### 3.1 Mutual information

The maximum uncertainty for a variable with *N* possible states is *H*_max_ = log(*N*) [36] and we can define the information *I* of a system simply as

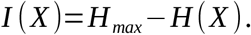

It is also possible to measure the uncertainty of *X* conditioned on another random variable *Y*

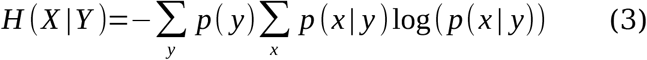

and then the mutual information between *X* and *Y* is the reduction in uncertainty about *X* thanks to the knowledge we have about *Y*

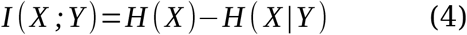

### 3.2 Relative entropy or Kullback-Leibler divergence

Kullback-Leibler divergence (*D*_KL_) or relative entropy allows us to compare two distributions *P* and *Q*, and quantifies the information gained by using *P* instead of *Q*.

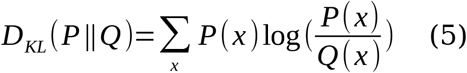

it can be shown (information inequality theorem) that *D*_KL_ ≥ 0, the equality being fullfilled only when *P* and *Q* are equal.

The relationship of *D*_KL_ with mutual information is

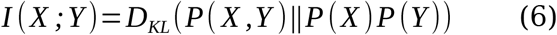

That is, mutual information quantifies the information gained by using the joint distribution of *X* and *Y* instead of the product of the two marginal distributions.

We can also relate *D*_KL_ to information and maximum entropy since

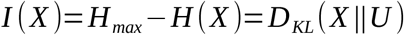

where *U* is the uniform over *X* [36].

### 3.3 Jeffreys divergency

Jeffreys divergence is the symmetric version of the *D*_KL_ divergence, that is, the sum of the two possible divergences between *P* and *Q* and therefore, it measures the total information that changes when using *Q* instead of *P* plus that of *P* instead of *Q*

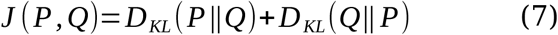

As a curiosity, *J* is also known as the Population Stability Index in the world of financial risk modeling and analysis.

Both *D*_KL_ and Jeffreys divergences are particular cases of the *f*-divergences which is defined, for the discrete case, as

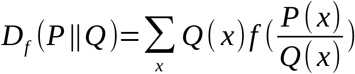

where *f* is the generating function which for *D*_KL_ is *f*(*x*) *= x*log *x* and for *J* is *f*(*x*) = (*x*-1)log *x*.

## 4. INFORMATION INTERPRETATION OF THE PRICE EQUATION

The first connection between the Price equation and information metrics is established by Frank [17] connecting the Price equation with Fisher information. The relationship of the Price equation with relative entropy and Jeffreys (*J*) divergence is made later by this same author [18,19,37,38].

To see the connection with the *J* divergence, let us return to equation (1) focusing on the first term which expresses the average change in the character z caused by selection [18]

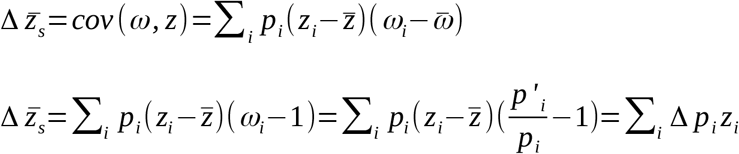

note that relative fitness is *ω*_*i*_ = *p’*_*i*_/*p*_*i*_, where *p*_*i*_ is the frequency of class *i* before selection and *p’*_*i*_ is the frequency after selection. The average relative fitness is equal to 1, Δ*p*_*i*_ = *p’*_*i*_ - *p*_*i*_ and ΣΔ*p*_*i*_=0.

If we take as character *z* the logarithm of the relative fitness, we obtain the expression of the first component of the Price equation in terms of information [18]

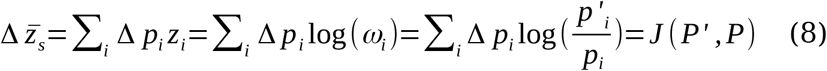

We see then that the mean change in the logarithm of relative fitness is the Jeffreys divergence between the frequency distribution after selection (*P*’) and before selection (*P*).

It is interesting to note that

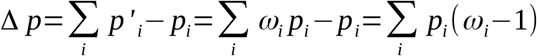

and

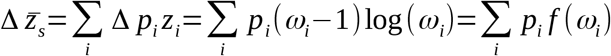

where *f* is the *f*-divergence generating function for *J*, so Eq. (8) is an *f*-divergence with generating function (ω - 1)log ω which explains that the evolution of the distribution caused by the fitness parameter in replicator-type models can always be expressed as a Jeffreys divergence.

Recently, Carvajal-Rodríguez uses these ideas to reinterpret Price’s equation in terms of the comparison between two sets of pairs, one representing random mating (*Q*) and the other (*Q*’) representing mating according to mutual mating fitness [23–25,39].

## 5. INFORMATION AND NON-RANDOM MATING

Sexual selection, defined as any type of selection arising from differential fitness in access to gametes for fertilization [40], can essentially be driven by two biological processes involving non-random mating: competition for mates and mate choice [25 and references therein].

In previous work, already mentioned, Carvajal-Rodríguez proposed a simple and abstract framework to describe mate competition and mate choice, along with their consequences in terms of sexual selection and assortative mating. This model is based on the concept of mutual and individual mating fitness. The effects of variation in these fitness measures are described in terms of sexual selection and assortative mating, quantified by the increase in information gained compared to random mating (zero information).

In what follows we will review how the informational interpretation of the Price equation connects with the information generated by non-random mating.

Let *q*’(*x*) be the frequency of females in matings, previously noted as *p’*_1_in [23], and *q*′(*y*) the frequency of males in matings (previously noted as *p*’_2_). Similarly, let *q*(*x*) be the population frequency of females (previously *p*_1_) and *q*(*y*) the frequency of males (previously *p*_2_). The joint distribution of random matings is *q*(*x,y*), where *q*(*x,y*) = *q*(*x*)*q*(*y*). Then, the joint distribution *q*’(*x,y*) according to the mutual mating fitness will be equal to [23]

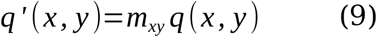

Model (9) relates the frequency *q*’(*x,y*) of matings guided by the mutual fitness *m*_*xy*_ for mating between females of class *x* and males of class *y*, with the frequency *q*(*x,y*) = *q*(*x*)*q*(*y*) of random mating between those female and males.

For each pair of classes we can define a feature or property *z*(*x,y*) about those classes such that the Price equation for the mean change in *z* caused by the variation in the mutual propensities *m* would be

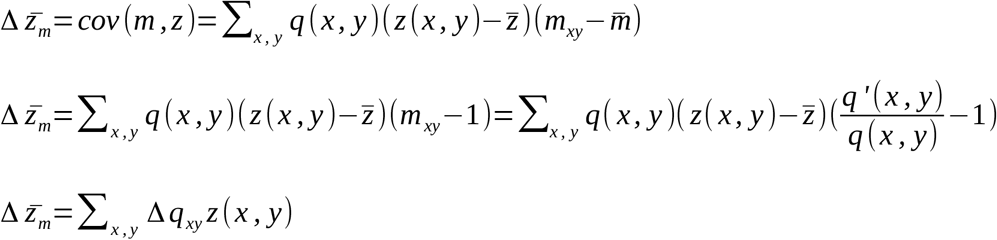

where Δ*q*_*xy*_=*q*’(*x,y*)-*q*(*x,y*) and ΣΔ*q*_*xy*_=0.

So, taking *z*(*x,y*)=log(*m*_*xy*_)=log(*q*’(*x,y*)/*q*(*x,y*)) we have that

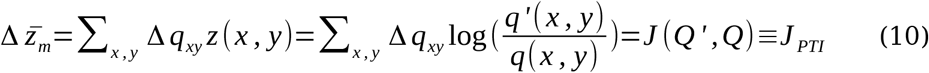

as obtained in [23].

Since *q*’(*x,y*)/*q*(*x,y*) when calculated on observed frequencies coincides with the pair total index (*PTI*) definided in [41], *J*(*Q*’,*Q*) has previously been called *J*_*PTI*_ and here we will maintain that notation, so *J*(*Q*’,*Q*) ≡ *J*_*PTI*_.

The *J*_*PTI*_ divergence in (10) can be decomposed into the sum of the information associated with sexual selection (*J*_*PSS*_), plus the information from assortative mating (*J*_*PSI*_), plus an interaction that arises when both, sexual selection and assortative mating, are present [23,25]. However, when the covariance within pairs is zero, *J*_*PTI*_ reduces to the information corresponding to sexual selection, denoted as *J*_*PSS*_ (see below). When the covariance within pairs is not zero, but the mean and variance between matings and the population are the same, *J*_*PTI*_ reduces to the information corresponding to assortative mating, referred to as *J*_*PSI [25]*_.

### 5.1 Information and sexual selection

As we have said, *J*_*PSS*_ information can be obtained from *J*_*PTI*_ when the covariance in the pairings is 0. This information is the sexual selection information, *J*_*PSS*_, and it compares the frequencies in the pairings *q*’(*x*)*q*’(*y*) with the population frequencies *q*(*x*)*q*(*y*). In [23] it is proved for the discrete case, that *J*_*PSS*_ is, in turn, the sum of the sexual selection information within each sex, *J*_*PSS*_ = *J*_*S1*_+*J*_*S2*_, and in [25] the same is proven for the continuous case. Here we will give another demonstration for both cases, continuous and discrete, based on the chain rule for relative entropy *D*_*KL*_ [36].

We know that

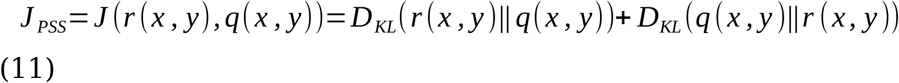

where *r* (*x, y*)=*q’* (*x*)×*q’* (*y*) and *q* (*x, y*)=*q* (*x*)×*q*(*y*).

And considering each sex separately

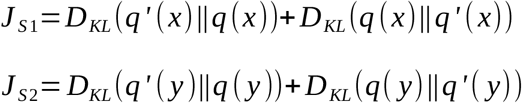

We want to show that *J*_*PSS*_ = *J*_*S1*_+*J*_*S2*_. To do this, let us consider the conditional probabilites

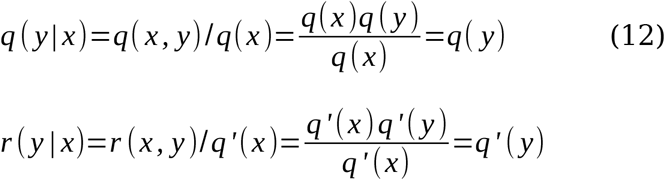

By the chain rule [36] we know that

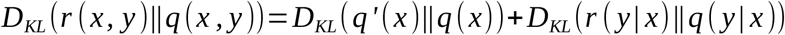

which due to (12) is

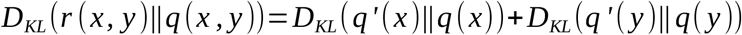

Similarly

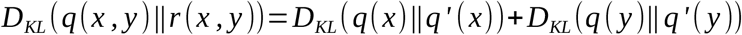

Therefore, if we add both divergences and group the *x* and *y* components separately we see that

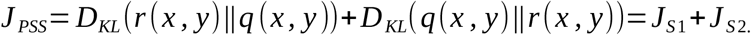

#### 5.1.2 Sexual selection in normally distibuted traits

If traits *x* and *y* are normally distributed, such that mating frequencies, *q*’, follow an *N*(μ_1_, Σ_1_), and population frequencies, *q*, follow an *N*(μ_2_, Σ_2_) where Σ_i_ is the variance-covariance matrix; then assuming that phenotypic changes are gradual [25] the components *J*_*S*1_ and *J*_*S*2_ coincide with the opportunity for sexual selection index [42].

On the other hand, if we consider the case where the variance in mating is equal to the population variance σ_1xx_=σ_2xx_ = σ_xx_ and σ_1yy_=σ_2yy_= σ_yy_, we have that *J*_*S*1_ and *J*_*S*2_ equal the square of the standardized sexual selection indices of females (*x*) and males (*y*), respectively. Furthermore, in this case, if there is no assortative mating, the relative entropy is symmetric so that *D*_KL_(*q*’||*q*) = *D*_KL_(*q*||*q*’) such that *J*_*PSS*_ = 2*D*_KL_(*q*’||*q*). Therefore, when the trait mean is the only difference between matings and the population, the sum of the squares for both sexes of the standardized sexual selection indices is equal to twice the relative entropy between mating frequencies and population frequencies.

To demonstrate the above, we first obtain *J*_*PSS*_ assuming that both traits are normally distributed. To do so, we note that the relative entropy between the *Q*’ distribution of expected pairings based on mutual fitness, and the *Q* distribution of expected pairings by chance is [25]

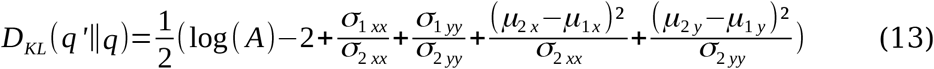

Where

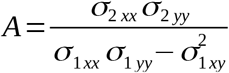

where it has been taken into account that the population covariance in the case of random encounters is 0 i.e., σ_2xy_=0.

The other relative entropy is

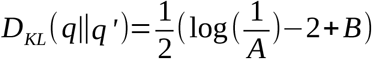

where

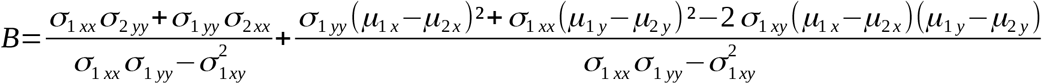

We have already seen that if there is no assortative mating *q*’(*x,y*) = *q*’(*x*)*q*’(*y*) = *r*(*x,y*) and the covariance within matings is zero, σ_1xy_=0, then

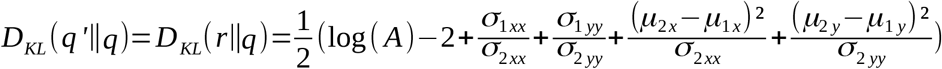

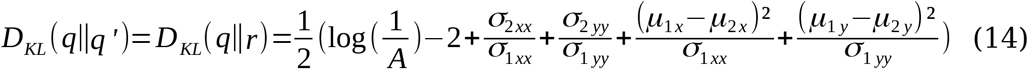

and

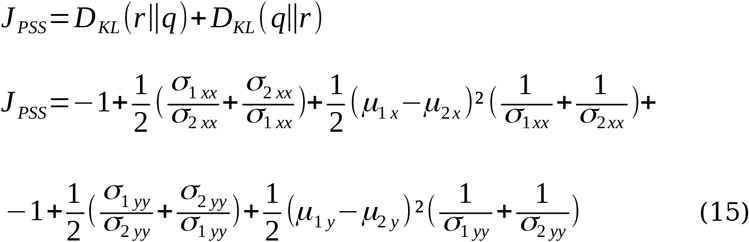

where we have regrouped the terms with *x* and *y* so that they correspond to *J*_*S*1_ + *J*_*S*2_.

Now, in the case where σ_1xx_=σ_2xx_ = σ_xx_ and σ_1yy_=σ_2y y_= σ_yy_, we see that by substituting in Eq. (14) we obtain *D*_KL_(*q*’||*q*) = *D*_KL_(*q*||*q*’) and by substituting in Eq. (15) we obtain *J*_*PSS*_ = *SSI*^2^_1_+*SSI*^2^_2_ where *SSI*_*1*_ and *SSI*_*2*_ are the standardized sexual selection indices for females and males, respectively, and therefore we have that

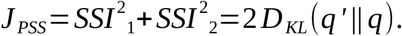

That is, the information on sexual selection, when there are no covariances and the variances do not change in the matings with respect to the population, is the sum of the square of the standardized selection indices, which in turn coincides with twice the relative entropy between the mating frequencies and the population frequencies.

### 5.2 Information and assortative mating

The *J*_*PTI*_ divergence, when we ignore differences between mating and population frequencies and focus only on frequencies within matings, becomes the *J*_*PSI*_ divergence which compares, within matings, the distribution due to mutual mating fitness with random mating

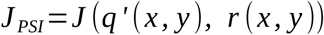

In previous work, *J*_*PSI*_ has been solved for both the discrete case [23] and the continuos case [25] based on the concept of mutual mating fitness along with the informational interpretation of Price’s equation. Here, for the continuous case and assuming a normal distribution, we will derive *J*_*PSI*_ directly from the sum of the relative entropy values when the mean and variance between the mating sample and the population are equal, and will see that it corresponds to a function of the square of the correlation ρ, which is interesting because ρ is typically used to detect assortative mating, but here we will show that the information corresponding to that pattern is actually captured by ρ^2^ / (1 - ρ^2^).

As before, we assume that *x* and *y* are normally distributed, such that *q*’ is *N*(μ_1_, Σ_1_) and *q N*(μ_2_, Σ_2_) where Σ_i_ is the variance-covariance matrix. As we have seen, the *J*_*PSI*_ divergence focuses only on the matings and the comparison within matings involves two groups of equal mean and variance for the trait, so we assume μ_1_ = μ_2_, σ_1xx_ = σ_2xx_ = σ_xx_ and σ_1yy_ = σ_2yy_ = σ_yy_, what will vary is the covariance within matigns σ_1xy_.

Then, from Eq. (13) we get

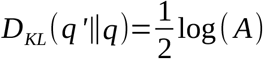

and the other divergence

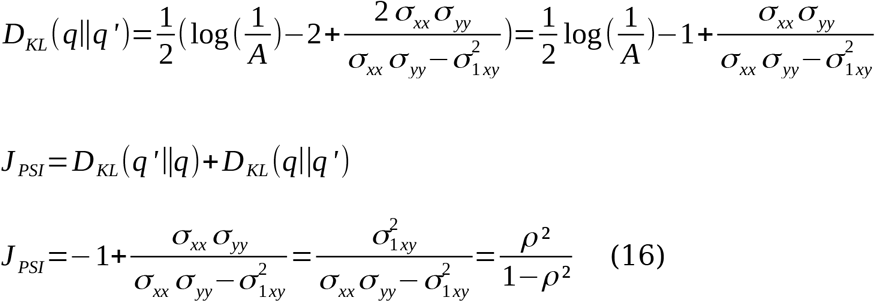

we see that assuming normal distribution in the matings, the assortative mating information is a function of the squared correlation coefficient [25].

## 6. INFORMATION AND EVOLUTION OF GENE FREQUENCY UNDER A GENERAL MATING AND FITNESS SCHEME

We have seen that the flow of information generated by the deviation from random mating is calculated by comparing the expected distribution according to mating fitness with the expected distribution of random matings. This information is obtained in the form of *J*_*PTI*_ divergence, regardless of whether the distribution is discrete or continuous, from which appropriate statistics are generated for each case [23,25]. However, the model (9) from which we derived *J*_*PTI*_ in (10), describes phenotypic frequencies (or classes in an infinitesimal interval in the continuous case), without making any assumptions about the underlying genetic model or connecting with the biological fitness of the offspring from those matings.

The aim of this section is precisely to connect the description of the mating distribution with the description of the evolution of gene frequencies subject to natural selection and to quantify all of this in terms of information. That is, to generate a model of gene frequency change for any explicit mating model and multilocus diploid genetic system and describe it in terms of information. This remains a pending task for classical population genetics and speciation models [43], although recently, information-based models have been developed that transcend the limitations of previous ones. See, for example [20] and [22]. These studies use *D*_KL_ as a metric that captures the information due to natural selection. However, the distributions these authors use are different, as Smith calculates the information associated with the trajectories by jointly considering the transitions and their states. In any case, the choice of *D*_KL_ is ad hoc in the sense that it is defined a priori to compare distributions and does not arise naturally from a model of change due to selection, as *J*-divergence does in the case of [18] or, in the case of deviation from random mating and sexual selection [23,25].

In what follows, I will propose a model that explicitly incorporates the mating system (the interindividual mating interaction) and implicitly the genomic context (mutation, linkage, etc.) into the classical replicator system. From this model, we see that the mean change in log-fitness caused by the change, at different stages, in the distributions of the mating system, genotypes and haplotypes, can be captured as information in the form of Jeffreys divergence.

We will solve the model for the simple case of one locus, 1) with random mating, thereby recovering the classical replicator system, and 2) with non-random mating but neutral in terms of viability, thus recovering the mating fitness model defined in [23]. In this last case, we will use as an example a simple one-locus hybrid sterility model to compare the information of the different Jeffreys divergences obtained.

### 6.1 General model

Let us consider a haplotype *H* of known frequency in generation *t, f*(*H*)_*t*_, as well as the frequencies of the corresponding diploid genotypes *HH, Hh* and *hh* where *h* represents any haplotype other than *H*. Likewise the joint distribution *q*’(*x,y*) according to the mutual mating fitness for the classes *HH, Hh* and *hh* would be given by (9)

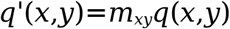

where *x* represents any female belonging to any of the three classes and *y* represents a male. Then, the general equation for the frequency of *H* in the next generation would be

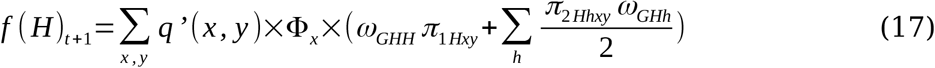

Eq. (17) connects the haplotype frequency *f*(*H*) in the next generation with the distribution of matings *q*’(*x,y*), the fecundity ć_x_ of female *x*, and the *G* genotypes of the offspring, carriers of the haplotype. Of these genotypes a proportion π_1_ will be homozygous and π_2_ heterozygous. The genotypes will survive based on their relative fitness ω (viability) after the action of natural selection.

The equation is general in the sense that the haplotype component *H* incorporates any multilocus genetic system, and the *q*’ component can incorporate any mating scheme. However, it is true that the values of π will depend on mutation and linkage disequilibrium, which in turn depends on the recombination frequency. Therefore, the model will not be dynamically sufficient unless we specify functions for the proportions π, which will depend on the case under study. In each case, we will obtain new statistics for detecting sexual selection and assortative mating, as well as new insights into the underlying evolutionary dynamics.

If there are only two types of haplotypes, the above formula can be simplified to

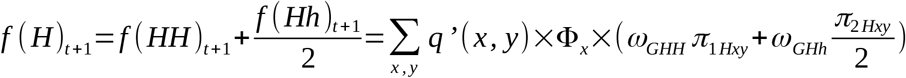

We will now apply (17) to the simple case of a biallelic locus without mutation.

### 6.2 One locus model

Let us consider a biallelic locus that codes for a discrete trait with 3 phenotypes, which are directly associated with their corresponding genotypes and follow a mating scheme described by the mutual mating fitnesses *m*_*xy*_. Within these mating fitnesses we also include the fecundity component *Φ*. We wish to determine the frequency *f*(*A*) of the allele *A* after mating, fecundity, and viability selection. The frequency *f*(*A*) after selection will be the sum of the frequency of the new *AA* genotypes plus half of the *Aa* genotypes. The structure of the model is as follows:

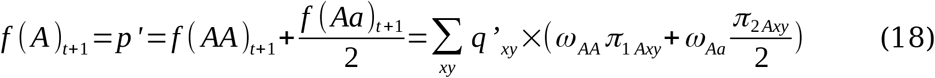

for convenience, we denoted *q*’(*x,y*) as *q*’_*xy*_, which therefore corresponds to equation (9). π_1*Axy*_ is the fraction of offspring from the mating *x*×*y* that are homozygous for the allele *A*, π_2*Axy*_ is the fraction of offspring from the mating *x*×*y* that are heterozygous for the allele *A, ω*_*AA*_ is the relative viability (divided by the average fitness) associated with the homozygote *AA*, and *ω*_*Aa*_ is the relative viability associated with the heterozygote.

To simplify the notation concerning pair parameters, we will use the subscript 1 for the *AA* genotype, 2 for the heterozygote, and 3 for *aa*. Thus, the genotype frequencies in the population before mating are *p*_1_ (*AA* frequency), *p*_2_ (*Aa* frequency), and *p*_3_ (*aa* frequency). If mating is random and we assume that there is no selection for fecundty, i.e., we take values *m*_*xy*_=1 for all *x, y*; the proportion of *AA* and *Aa* genotypes in the offspring before selection and without mutation would be as shown in Table 2.

**Table 1.** Table of main symbols used in section 6.

| Symbol | Meaning |
| --- | --- |
| $q$ | mating frequency if mating is random. |
| $m_{xy}$ | relative mutual mating fitness between female with phenotype $x$ and male with phenotype $y$, i.e. the propensity of mating with fertilization between a given pair of phenotypic classes. |
| $q'$ | mating frequency after the action of mutual mating fitness. |
| $\Phi_G$ | fecundity, it is included within the mutual mating fitness. |
| $M$ | average mutual mating fitness. |
| $m_G$ | the relative (divided by the average $M$) marginal mating fitness of genotype $G$. |
| $\omega_G$ | the relative (divided by the average fitness $W$) genotype viability i.e. the potential for survival to adult of the given genotype. |
| $\Omega_{xyG}$ | the product of $m_{xy} \Phi_G \omega_G$. |
| $H$ | a given haplotype i.e. a sequence that carries a particular set of genes that tend to be inherited together. |
| $\pi_{1Hxy}$ | fraction of the offspring of the mating $(x, y)$ that are homozygous for the $H$ haplotype. |
| $\pi_{2Hxy}$ | fraction of the offspring of the mating $(x, y)$ that are heterozygous for the $H$ haplotype. |
| $p_1, p_2, p_3$ | For one locus, frequencies of the genotypes $AA, Aa$ and $aa$, respectively. These symbols are used to simplify the notation of mating frequencies in single-locus examples. |

**Table 2.** Proportion *p*_x_*p*_y_π of offspring genotypes *AA* and *Aa* as a function of mating frequencies when *m*_*xy*_ = 1 for every *x, y*.

| Matings | Offspring |  |
| --- | --- | --- |
| | $\pi_{1A}$ (AA ) | $\pi_{2A}$ (Aa ) |
| $p_1 \times p_1 \times m'_{11} / M$ | $p_1^2$ | 0 |
| $p_1 \times p_2 \times m'_{12} / M$ | $(p_1 p_2) / 2$ | $(p_1 p_2) / 2$ |
| $p_1 \times p_3 \times m'_{13}/M$ | 0 | $p_1 p_3$ |
| $p_2 \times p_1 \times m'_{21}/M$ | $(p_1 p_2)/2$ | $(p_1 p_2)/2$ |
| $p_2 \times p_2 \times m'_{22}/M$ | $(p_2^2)/4$ | $(p_2^2)/2$ |
| $p_2 \times p_3 \times m'_{23}/M$ | 0 | $(p_2 p_3)/2$ |
| $p_3 \times p_1 \times m'_{31}/M$ | 0 | $p_1 p_3$ |
| $p_3 \times p_2 \times m'_{32}/M$ | 0 | $(p_2 p_3)/2$ |
| $p_3 \times p_3 \times m'_{33}/M$ | 0 | 0 |

Note that $m_{xy} = m'_{xy}/M$

If we assume random mating, by (9) we know that the probability of mating between *x* and *y* is *q*’_*xy*_ = *q(*x,y) = *p*_x_*p*_y_ and, noting the allele frequencies of the previous generation as *p* and *q* (without subscripts), we would have that *p*_1_=*p*^2^, *p*_2_=2*pq* and *p*_3_=*q*^2^. Therefore, substituting the values of Table 2 in Eq. (18) would be

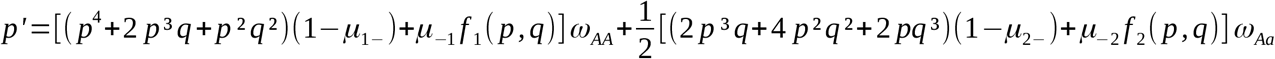

where μ__1_ and μ__2_ are any mutations producing genotype *AA* or *Aa*, respectively, μ_1__ is any mutation from *AA* to any other genotype and μ_2__ from Aa to any other genotype and *f*_1_ and *f*_2_ are functions of allele frequencies.

Assuming there is no mutation μ_,_ = 0 and after simplifying we obtain

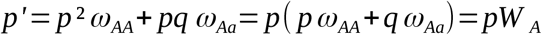

where *W*_A_ = *pω*_*AA*_+*qω*_*Aa*_ is the marginal fitness of allele *A* [44].

We see that, for a biallelic locus and with random mating, equations (17-18) correspond to the basic model of selection for a biallelic locus [44,45].

#### 6.2.1 Genotypic mating fitness

Alternatively, if we assume a neutral model such that *ω*_AA_ = *ω*_Aa_ = *ω*_aa_ = 1, Eq. (18) becomes

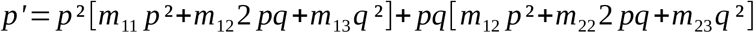

where we have assumed that the mutual mating fitnesses between classes *x* and *y* do not depend on the sex of the genotype, that is, *m*_xy_ = *m*_yx_ and that the genotypic frequencies are equal in males and females.

Under these conditions, the individual mating fitness [formula 4 in 24] for type *AA* females is

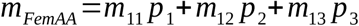

and for males,

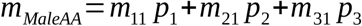

and since *m*_xy_= *m*_yx_ we have that *m*_FemAA_ = *m*_MaleAA_ = *m*_AA_ being

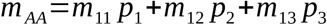

In this context, where each phenotypic class corresponds to a genotype, the individual mating fitness *m*_AA_ corresponds to the genotypic mating fitness for the *AA* genotype. Then, the frequency of the *AA* genotype in matings is

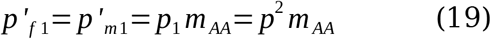

Similarly, for heterozygotes the genotypic mating fitness *m*_Aa_ is

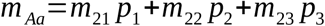

being the frequency of *Aa* in matings

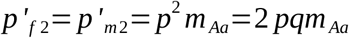

Therefore, under non-random mating selectively neutral model, the equation for the new frequency of allele *A* in the next genration can be rewritten as

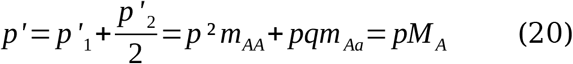

where *M*_A_ is the marginal mating fitness of allele *A*.

The frequency of allele *A* in the new generation is the sum of the frequencies of genotype *AA* plus half the frequency of the heterozygous genotype *Aa* after mating selection. Note that if the genotypic mating fitnesses are 1, meaning that mating is random, then *p*’ = *p*, as expected.

Similarly, for the genotype *aa* we have that

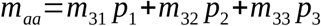

then *p*’_3_ = *q*^2^*m*_aa_ and the frequency of the allele *a* in the next generation is

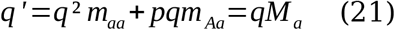

where *M*_*a*_ is the marginal mating fitness of allele *a*.

Finally, the change in gene frequency due to differential mating fitness will be

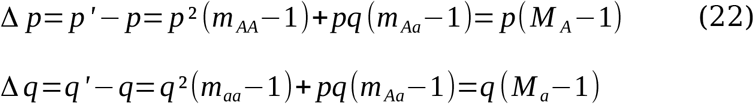

The sum of frequencies *p*’+*q*’=*p*’_1_+*p*’_2_+*p*’_3_ must equal 1. To check this, remember that the mutual mating fitnesses *m*_*xy*_= *m*’_*xy*_/*M* are normalized (Table 2) and therefore the genotypic fitnesses are also, *m*_AA_=*m’*_AA_/*M, m*_Aa_=*m’*_Aa_/*M* and *m*_aa_=*m’*_aa_/*M*, where the tilde is used to indicate the unnormalized fitness. The mean fitness *M*=Σ_xy_ *p*_x_*p*_y_*m*’_xy_ can be expressed in terms of the unnormalized genotypic fitnesses as

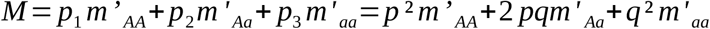

Therefore using (19-21) the sum *p*’+*q*’ = (*p*^2^*m*’_AA_ +2*pqm*’_Aa_+*q*^2^*m*’_aa_)/*M* = *M*/*M* = 1.

### 6.3 Information and fitness

Eq. (17) can be expressed in term of diploid genotype frequencies and a combined fitness by substituting with Eq. (9) *q*’_*xy*_=*m*_*xy*_*q*_*xy*_, we obtain

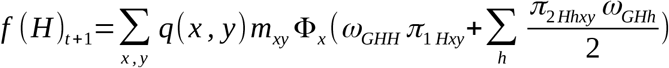

therefore, since the haplotype frequency is simply the sum of the frequencies of the genotypes containing that haplotype, we can also express the frequency of any genotype *G* as

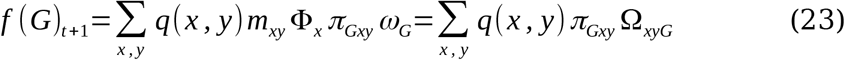

where Ω_xyG_ = *m*_*xy*_ć_x_*ω*_*G*_.

The genotype frequencies in the previous generation can be expressed as

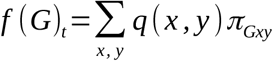

For example, if *G* = *AA* then *f*(*AA*)_*t*_ is the sum of the expected frequency by chance of crosses *AA* **×** *AA, Aa* **×** *Aa, AA* **×** *Aa* and *Aa* **×** *AA* and considering the corresponding proportion of *AA* genotype in the offspring of each cross, i.e.

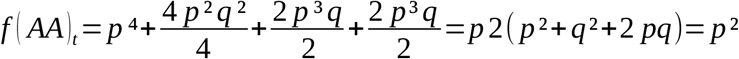

Note that in general the overall fitness component of *G*, Ω_G_, is different from the frequency ratio before and after selection

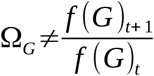

because for each mating we have Ω_xyG_ = *m*_*xy*_ć_x_*ω*_*G*_ that can differ for differents matings *x, y*.

The only fitness element common to genotype *G* will be viability ω_G_, so if we assume no differences in mating fitness or fertility, we have that

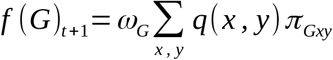

So

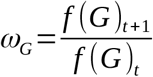

and in this later case, the mean change in log-fitness caused by selection due to differential viability log-*ω*_*G*_ is

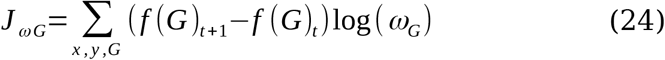

#### 6.3.1 Information on matings by genotype

However, we are interested in the information due to changes in genotypic frequencies when the different fitnesses causing that change may vary. The problem is that the final frequency of each genotype, which is due to the different mating fitness of its parents, the fecundity of the mother, and ultimately the viability of the genotype itself, cannot be expressed as the result of a single measure of fitness.

One option will be to obtain the information associated with each genotype separately. That is, the average change in the logarithm of the overall genotypic fitness, Ω_*xyG*_, after mating (*x, y*) associated with the change in frequency of that genotype from one generation to the next is

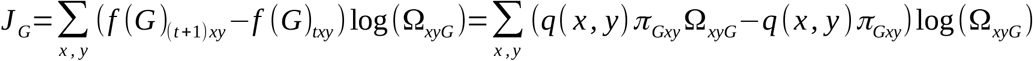

where note that

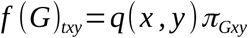

Therefore, for the entire set of possible genotypes *G* whose frequency changes from one generation to the next due to the effect of selection, we would have

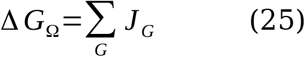

That is, if the combined effect of mating fitness, fecundity and viability produces a shift in genotype frequencies, this can be captured as sum of Jeffreys divergences.

#### 6.3.2 Genotype distribution information

Alternatively and from Eq. 23 we can directly consider the frecuency of a genotype as

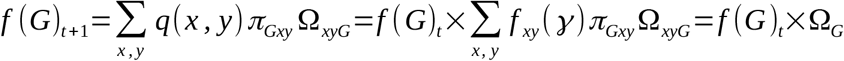

where

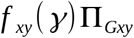

is a functional of the genotypes frequencies after the (*x, y*) mating, and Ω_*G*_, is the overall marginal fitness of genotype *G*

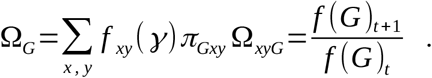

So, the mean change in this overall log-fitness caused by the variation in genotype frequecies because of selection takes the form of Jeffreys divergence as

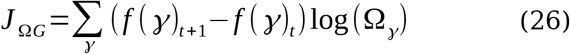

#### 6.3.3 Haplotype distribution information

From Eq. 17 we can directly consider the frecuency of haplotype *H* as

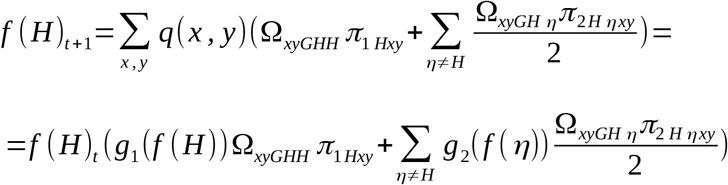

where *η* is any allele other than *H* and *g*_1_ and *g*_2_ are funcionals of the frequency in the previous generation of *H* and *η*, respectively.

Then

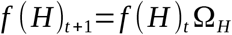

where

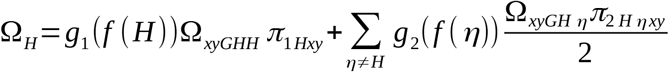

is the marginal overall fitness of haplotype *H*.

So, the total mean change in log-fitness caused by the change in haplotype frequencies is

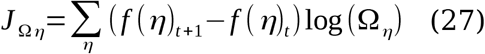

Therefore, from model (17), if we measure the mean change in log-fitness we can obtain different measures of information associated with the change in genotype frequency or the change in haplotype frequency. These types of information are generally different since there can be genotypic change without a change in gene frequency. As for the change in genotypes we calculate the information in two ways, one linked to the variation of each genotype obtained from different matings and another linked to the variation in the genotype frequencies as a whole in the new population.

Let’s see below some simple examples for a biallelic locus.

#### 6.3.4. Sexual selection information for a biallelic locus

We will assume as before, that the genotypic frequencies in males and females are equal, in the case of females we already saw that, *p*’_*f*1_ = *p*_*f*1_*m*_*AA*_, *p*_*f*1_ = *p*^2^, *p*’_*f*2_ = *p*_*f*2_*m*_*Aa*_, *p*_*f*2_ = 2*pq*, etc… The difference between the population frequencies *p*_*f*_ and the frequencies within the matings *p*’_*f*_ is Δ*p*_*f*1_=*p*’_*f*1_ - *p*_*f*1_ = (*p*_1_)*m*_AA_ -(*p*_1_), etc.

Information due to sexual selection in females in terms of genotypic mating fitness will be

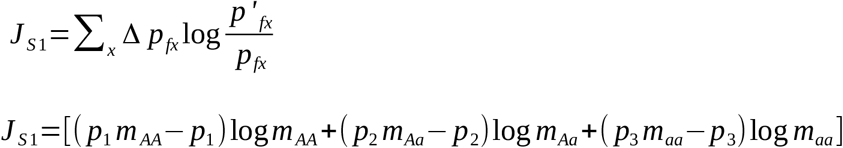

and since the genotypic frequencies in males and females are equal *p*_*fx*_ = *p*_*my*_ and we have assumed that the mutual fitnesses are symmetrical *m*_*xy*_ = *m*_*yx*_ then, sexual selection in males is equal to that of females *J*_*S*2_ = *J*_*S*1_ and since *J*_*PSS*_ = *J*_*S*2_+*J*_*S*1_ then

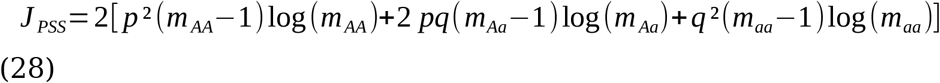

where we can see that Jeffreys’ divergence for sexual selection captures the information generated by the change in genotype frequencies weighted by the logarithm of the genotypic mating fitness.

We can now compare in this same scenario what the genotypic information (Eq. 25-26) and the haplotypic information (Eq. 27) would be.

The Eq. 25, computes the information generated by the change in the distribution of genotypes as a sum of informations

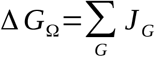

where

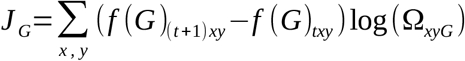

in our case this implies, recalling that *m*_*xy*_ = *m*_*yx*_ and that Ω_xyG_ = *m*_*xyG*_, then

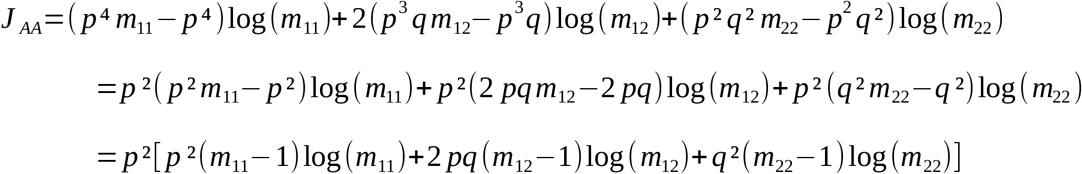

similarly,

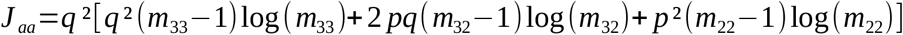

and finally

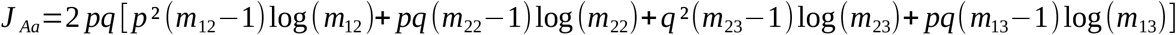

now recall that

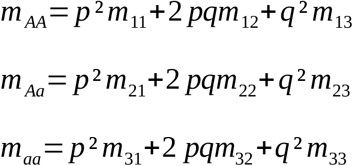

we see that the difference between *J*_*PSS*_ and *J*_*G*_ is where we focus the unit of information. That is, for the sexual selection information *J*_*PSS*_, the focus is in each genotype which is involved in various matings. So, for *m*_*AA*_ the focus is in the combination of matings that involve one or two *AA* genotypes as parents.

However, for the *J*_*G*_ information the focus is within a specific genotype produced by the matings.

As for the other measure of information that we have defined for the genotypes (Eq. 26)

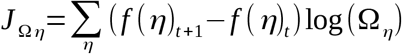

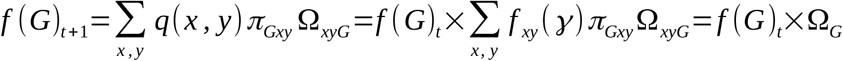

which in our sexual selection scenario implies

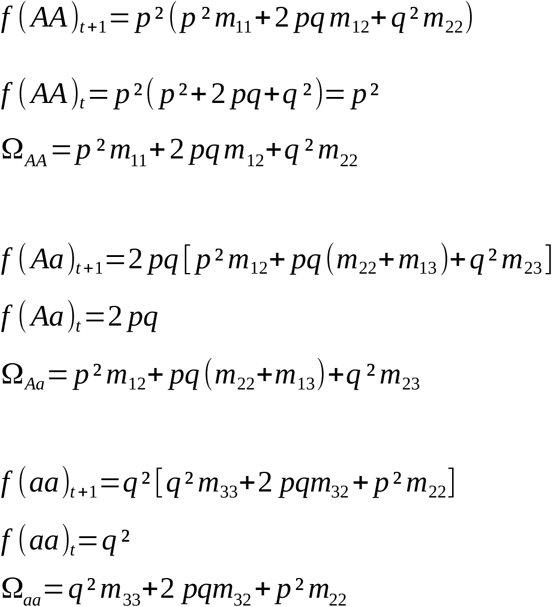

so that,

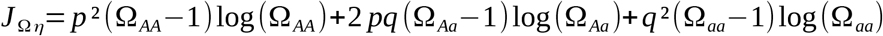

We see that the difference between *J*_*PSS*_ in Eq. 28 and *J*_*Ωη*_ is due to the different fitness weights of each genotype and, since *m*_*xy*_ = *m*_*yx*_, when comparing we see that

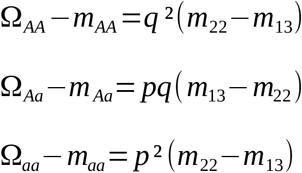

The difference lies in the crosses *Aa* **×** *Aa* (*m*_22_) and *AA* **×** *aa* (*m*_13_ or *m*_31_). This is because the *J*_*PSS*_ calculation of sexual selection focuses on the variation generated by matings on the marginal frequency of each genotype. For example, the marginal frequency of *AA* comes from the matings of *AA* with other genotypes, which includes the cross *AA* × *aa* (mating fitness *m*_13_) but not the *Aa* × *Aa* (mating fitness *m*_22_). The same applies to the marginal frequency of the aa genotype; it includes the cross *aa* × *AA* but not the cross between heterozygotes.

On the other hand, the calculation of the genotypic information *J*_*Ωη*_ directly studies the variation in the frequency of each genotype resulting from a set of matings. In this case, the pattern is reversed: for the genotypes *AA* and *aa*, the cross between heterozygotes is included, but not the *AA* × *aa*. The opposite holds true for the *Aa* genotype.

In this specific case of a biallelic locus with no differences in viability or fertility, but with mating fitness under the constraint *m*_*xy*_ = *m*_*yx*_, if the mating fitness among heterozygotes coincides with the mating fitness among different homozygotes, the information obtained from the genotypes aligns with the information from sexual selection. If this condition is not met, the information is similar but differs by a factor associated with the difference *m*_22_-*m*_13_.

Finally, for the haplotype information measure (Eq. 27), the information generated by the change in the distribution of haplotypes is

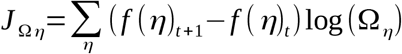

where

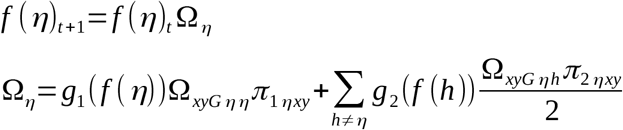

that in our case implies

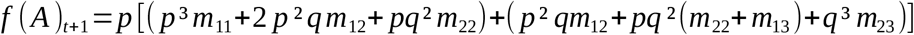

so

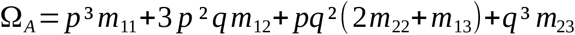

and

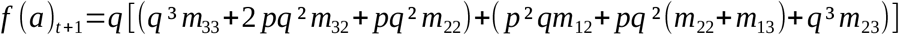

so

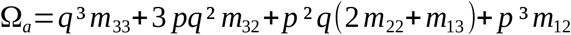

Information about the change in allele frequency caused by selection is

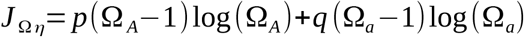

Note that if the mating fitnesses *m*_*ij*_ are equal to one, then Ω_A_ = Ω_a_ = 1 and therefore the information is zero.

#### 6.3.5 Toy example: Information for hybrid sterility

Let us compare the different information indices we have defined using a simple hybrid sterility model. We continue with the one locus model without mutation and assume that heterozygotes have lower fertility so that their frequency of successful mating (with fertilization) is lower. That is, *m*’_12_ = *m*’_21_ = *m*’_22_ = *m*’_23_ = *m*’_32_ = ε < 1. The rest of the mating fitnesses equal one, *m*’_11_ = *m*’_13_ = *m*’_31_ = *m*’_33_ = 1. The average mating fitness is

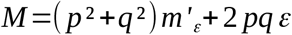

where

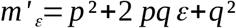

Since the frequencies of males and females are equal and *m*’_*ij*_ = *m*’_*ji*_, then the marginal fitnesses of females and males are equal

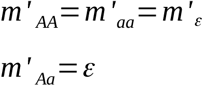

the normalized frequencies will be

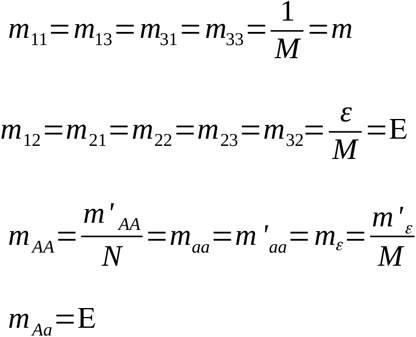

##### 6.3.5.1 Sexual selection information

Substituting in (28) we obtain

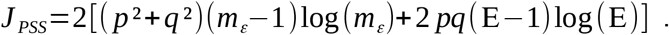

*J*_*PSS*_ represents the information obtained from the change in the phenotypic frequencies of each sex (in this case equal to the genotypic frequencies) within the matings with respect to the population frequencies.

Note that in the particular case of equal allelic frequency, *p* = *q*, we have that

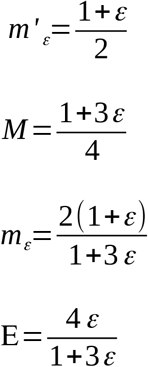

and the value of *J*_*PSS*_ is not 0,

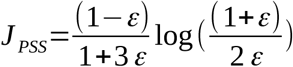

we see that if, for example, we take ε = 1/3, we obtain *J*_*PSS*_=(1/3)log(2) ≠ 0. This is interesting because we will see that this model with equal allele frequencies does not affect the distribution of genotypes in the following generation and therefore the information obtained from the distributions of genotypes and haplotypes (alleles) will be 0 (see below).

##### 6.3.5.2 Information on matings by genotype

We can calculate the sum of the *J*_*G*_ indices that for each genotype compute the frequency of the genotype after the matings with respect to the expected frequency if the matings are random. This mating information on each genotype corresponds to Eq. 25 and in our example is

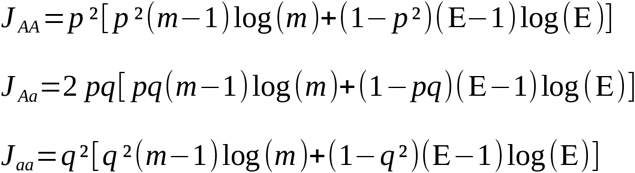

The total change for the three genotypes will be the sum

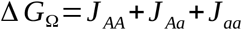

Note that in the case *p* = *q, J*_*AA*_ = *J*_*aa*_=*J, J*_*Aa*_ = 2*J*. Then Δ*G*_Ω_ = 4*J*.

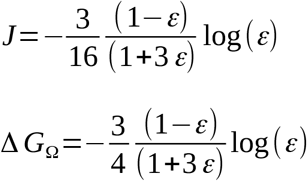

taking ε=1/3, Δ*G*_Ω_ = (1/4)log(3) ≠ 0.

We see that the mating information by genotype is not 0 either, even though the allele frequencies are equal. This is because the distribution of genotypes by mating changes, even though the total distribution remains constant.

##### 6.3.5.3 Genotype distribution information

We can use Eq. 26 to calculate the information obtained when the distribution of genotypes changes. In this case, the fitness associated with each genotype is

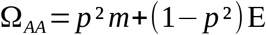

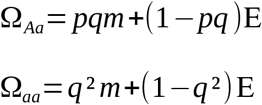

And the information is

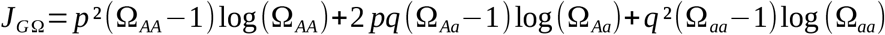

note that if *p* = *q* = 0.5 the three values of Ω_G_ are equal and the value *m* = 4/(1+3ε) and E =4ε/(1+3ε) so that

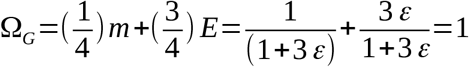

and the information *J*_*ΩG*_ = 0.

This is expected since selection acts on heterozygotes and the gain in frequency of homozygotes is distributed equally (in the case of *p* = *q*), generating heterozygotes again and therefore the genotypic distribution is not altered when the frequencies of both alleles are equal.

##### 6.3.5.4 Haplotype distribution information

We calculate it from Eq. 27 and we obtain

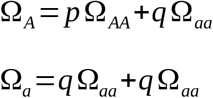

and the corresponding information is

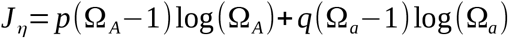

which, just like in the previous case, will be 0 if *p* = *q*.

## 7. DISCUSSION

In this work, we have proposed a population genetics model that begins with a mating system guided by mutual mating fitness between individuals, continues with the distribution of genotypes resulting from each mating, and follows with the subsequent survival of these genotypes based on their viability fitness until they reach adulthood. At each stage of the model, variation in the distribution of genotypes is captured in terms of information.

Population genetics models are fundamental both for testing proposed hypotheses, whether general or associated with a specific species and evolutionary scenario, and for deepening our understanding of the history of species and biodiversity on Earth. They also help reveal invariances and logical connections that, in turn, may suggest new questions and pose new hypotheses [46,47].

As mentioned in the introduction, information theory seems to be increasingly relevant in providing new descriptions of the evolutionary process, allowing us to explore certain aspects more deeply or facilitate the development of models and estimators for parameters of interest. Recent studies have shown how selection accumulates information in the genome [20] and also how the flow of information throughout the process of evolutionary change can capture different degrees of adaptation and help develop new statistics [22].

It should be noted that when we consider typical equations of classical population genetics, or the replicator equation as *p*’=Ω*p* where Ω is some kind of relative fitness, and we express the mean change in the logarithm of the fitness of the character, what we obtain is an equation of the type

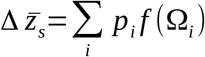

where *f* is the generating function (Ω - 1)log(Ω) of an *f*-divergence corresponding to the Jeffreys divergence.

Therefore, every component of the evolutionary process that is guided by some kind of selection, be it sexual selection, mate choice, viability, etc. such that we can express the expected change in terms of (Ω - 1)log(Ω) can be described in terms of the Jeffreys divergence. That is, the dynamics of evolutionary change itself, at least as it can be expressed in the given form *p*’=*p*Ω, can be measured in terms of the Jeffreys divergence, because that is what it is mathematically, in this sense it is not an ad hoc chosen statistic.

In the model presented here, the average change in fitness at different stages of the life cycle can be calculated as Jeffreys information. This has allowed us to calculate the information associated with various stages of the process, namely, the information obtained from the distribution of matings, the information associated with the matings fo each genotype, as well as the information associated with the distribution of genotypes and haplotypes in the new generation. We have found that these indices contain distinct information and therefore could allow us to draw inferences about associated phenomena.

Population genetics models typically assume random mating [20,46] or, at most, consider any mating system implicitly, as in [21,22], where different mating systems may be reflected in the topology of a graph. In our case, by explicitly modeling the mating system in terms of information, on the one hand, we unify different aspects related to the detection of sexual selection and assortative mating and facilitate the development of statistical measures [25]. And on the other hand, by incorporating mating fitness into a more general model, we reveal connections between different mating processes and other parts of the gene frequency evolution process, described in terms of information.

An interesting detail of the proposed model is that the average changes in different types of fitness, due to variation between the corresponding distributions, can be easily translated into statistical tests since the Jeffreys divergence approximates well a chi-square distribution when the hypothesis of equality between distributions is met [36,48].

All this has allowed the development of a multi-model selection methodology applied to the estimation of non-random mating [24] that can be used in empirical studies, see for example [49–51]

### 7.1 Limitations of the model

The proposed model connects the matings in the parental generation with the distribution of adult genotypes in the next generation. The model explicitly accounts for the mating system defined by mutual mating fitnesses, female fecundity, and the viability associated with genomic genotypes. The effects of mutation and recombination are not explicitly modeled, as they are implicitly included in the calculation of genotypic proportions following matings. However, in its current form, the model does not consider migration, genetic drift, epigenetic inheritance, or other potential factors related to niche construction or habitat choice [52].

However, these aspects are not central to the main message of the work, which emphasizes the connection between different aspects of evolutionary change and the measurement of information through shifts in distributions, a point also highlighted in recent studies [20,22].

## 8. CONCLUSION

In conclusion, it may be worth considering the relationship between biological fitness and information, whether the increase in information is favored by selection [53], or even if we can think of fitness in terms of a multilevel information alignment between populations and their environment [54,55]. Additionally, we might reflect on the advantages of shifting from a fitness-driven approach to evolution toward one guided by information and whether this could help clarify, broaden and deepen the mathematical description of evolutionary theory.

## Data Availability Statement

No data were used in this study.

